# Computational model for synthesizing auditory brainstem responses to assess neuronal alterations in aging and autistic animal models

**DOI:** 10.1101/2024.08.04.606499

**Authors:** Ben-Zheng Li, Shani Poleg, Matthew Ridenour, Daniel Tollin, Tim Lei, Achim Klug

**Affiliations:** Department of Physiology & Biophysics, University of Colorado School of Medicine, Aurora, Colorado, USA; Department of Electrical Engineering, University of Colorado Denver, Denver, Colorado, USA

**Keywords:** Computational model, auditory brainstem response, spiking neuronal network model, neural simulation, Fragile x syndrome, age-related hearing loss

## Abstract

**Purpose:** The auditory brainstem response (ABR) is a widely used objective electrophysiology measure for non-invasively assessing auditory function and neural activity in the auditory brainstem, but its ability to reflect detailed neuronal processing is limited due to the averaging nature of the electroencephalogram-type recordings.

**Method:** This study addresses this limitation by developing a computational model of the auditory brainstem which is capable of synthesizing ABR traces based on a large, population scale neural extrapolation of a spiking neuronal network of auditory brainstem circuitry. The model was able to recapitulate alterations in ABR waveform morphology that have been shown to be present in two medical conditions: animal models of autism and aging. Moreover, in both of these conditions, these ABR alterations are caused by known distinct changes in auditory brainstem physiology, and the model could recapitulate these changes.

**Results:** In the autism model, the simulation revealed myelin deficits and hyperexcitability, which caused a decreased wave III amplitude and a prolonged wave III-V interval, consistent with experimentally recorded ABRs in Fmr1-KO mice. For the aging condition, the model recapitulated ABRs recorded in aged gerbils and indicated a reduction in activity in the medial nucleus of the trapezoid body (MNTB), a finding validated by confocal imaging data.

**Conclusion:** These results demonstrate not only the model’s accuracy but also its capability of linking features of ABR morphology to underlying neuronal properties and suggesting follow-up physiological experiments.

## 1. Introduction

The auditory brainstem response (ABR) is a non-invasive or minimally invasive electrophysiological measurement of auditory evoked potentials created by auditory brainstem nuclei. To record ABRs, small, brief and in our case multifrequency sound stimuli (clicks) were presented to rodents. These clicks recruited simultaneous action potentials in auditory nerve fibers which can be measured noninvasively through the skull as wave I. As this activity ascends the lower auditory system, subsequent waves II to V are created, representing neural activity along the ascending auditory pathway [1]. This technique allows for simultaneous objective assessment of multiple stages of the auditory system, making it a useful tool for non-invasively detecting hearing impairments in clinical settings [2] as well as a frequently used tool in hearing research studies to investigate auditory dysfunction and underlying neurological disorders [3].

However, the ABR only measures the overall neural activity at each stage due to the use of low impedance electrodes which are placed outside the skull, similar in nature to EEG recordings which average across populations of neurons. Some stages, quantified as one single wave, may contain neural circuits with multiple nuclei and multiple steps of auditory information processing, such as signal sharpening, coincidence detection, and sound localization [4, 5]. These simultaneous activity patterns may not be independently reflected in ABR wave metrics and typically necessitate alternative methods of measurements such as patch-clamp, extracellular recordings or optogenetic manipulation [6–8]. Similarly, computational models for auditory information processing typically use parameters and measurements from single cell data [9, 10]. Validating model predictions involving the propagation and transformation of auditory information across multiple stations of the ascending auditory system may require recordings and interventions at multiple stages in a large population size, which is experimentally challenging and difficult to achieve at cellular level resolution.

Therefore, there is a need to model ABR waveforms based on underlying cellular properties, as well as a need to do the opposite - be able to predict changes in cellular physiology from alterations in global ABR morphologies. Several sophisticated ABR models are available to date. However, they are either based on population-level firing rates [11–13], lacking the ability to address detailed biophysical properties and membrane dynamics for single neurons, or they are based on simulating only a single stage of the auditory brainstem [14, 15]. Hence, there is a need for a computational model that involves the entire ascending neural circuitry of the auditory brainstem and simulates the complete ABR trace.

In this study, we developed a computational model of the auditory brainstem that can perform large-scale neural simulations of the afferent auditory pathway and synthesize ABRs I through V based on simulated neuronal activities evoked by modeled sound wave stimulation. We validated this model by computing the ABR of an autistic (Fragile X) animal line based on previously reported alterations and myelin deficits in Fragile X. We observed morphologically altered ABRs with decreased wave III and prolonged wave III-V, which were then validated with experimentally recorded ABRs from *Fmr1*-KO mice versus wild-type controls. Furthermore, this model was also used to predict alterations in the ABR recorded from aging gerbils, revealing a decreased population size in the medial nucleus of the trapezoid body (MNTB). This latter finding was experimentally verified by counting the number of MNTB cells in confocal image datasets captured from brain sections of aging gerbils.

## 2. Materials and Methods

### 2.1. ABR acquisition

The ABR recording procedures are similar to the approaches described in previous publications [16–18] and illustrated in Fig.1. ABRs were measured from *Fmr1*-KO mice (N = 22, 13 males and 9 females) and C57BL/6 wildtype mice (N = 20, 12 males and 8 females) with an age range of 73.03 ± 11.64 (Mean ± SD) days old. Animals were anaesthetized with a mixture of ketamine and xylazine by intraperitoneal injections. Platinum needle electrodes were placed under the skin at the apex (recording site), nape (reference), and back leg (ground) after the toe-pinch reflex disappeared. Sound stimuli were 0.2-ms complex clicks with 35 ± 5 ms (Mean ± SD) interstimulus intervals and were delivered by a pair of speakers (MF-1 multi-field speakers, Tucker Davis Technologies, Alachua, FL, USA) which received amplified soundwaves generated by a customized Python script. Recorded traces were filtered using a 50-3000 Hz second-order bandpass Butterworth filter and averaged across 1000 repetitions per condition, i.e., binaural, left monaural, and right monaural. In addition, the binaural hearing and spatial hearing were also assessed by the binaural interaction component (BIC) of the ABR. The BIC was calculated by taking the difference between the sum of the two (left plus right ear) monaural ABRs minus the ABR recorded with binaural sound stimulation on a point-by-point basis [17, 19]. All experimental procedures were reviewed and approved by the institutional animal care and use committees (IACUC) of the University of Colorado Anschutz Medical Campus (permit no.00617) and complied with all National Institutes of Health (NIH) and OLAW guidelines for the humane treatment of laboratory animals.

**Figure 1.**
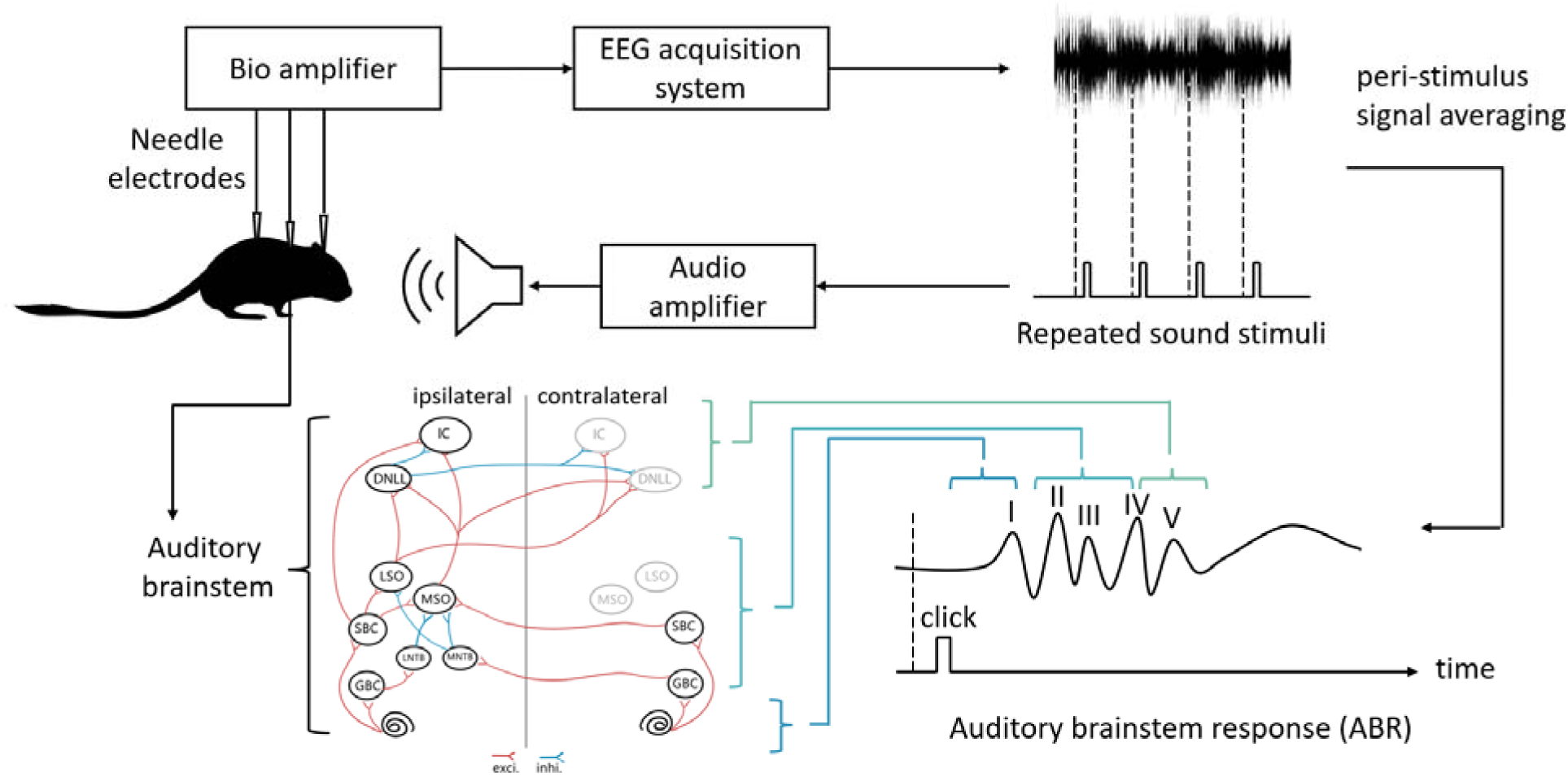
Schematic of the ABR recording setup (top) and sources of ABR waves (bottom).

### 2.2. Neuron and synapse models

Spiking neurons are the most basic building block to construct the expected auditory brainstem model. Multiple spiking neuron models are available, ranging from the leaky integrate-and-fire model, the Izhikevich model [20], up to much more biophysically realistic (and computationally demanding) Hodgkin-Huxley [21] and multicompartment models. Among these options, the conductance-based leaky integrate-and-fire model stands out as both computationally efficient and widely used, offering a strong balance of simplicity and biological plausibility. The dynamic of the membrane potential *v_m_* is described by the following equation:

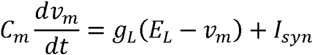

where *v_m_* is the membrane potential, (*C_m_* is the membrane capacitance, *I_syn_* is the synaptic current, *g_L_*, is the leaky conductance, and *E_L_*, is the leaky reversal potential. The leaky integrate-and-fire neuron emits a spike when its membrane potential reaches the firing threshold *v_th_*, and the membrane potential resets to *v_reset_* subsequently.

Synapses can be either simulated by a conductance-based model, where the synaptic current is given by the product of the conductance and the driving force,

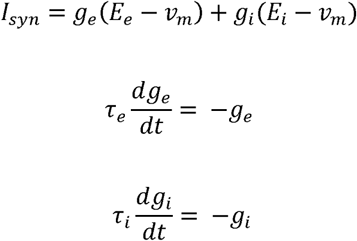

where *g_e_* is the excitatory conductance, *g_i_* is the inhibitory conductance, *E_e_* is the reversal potential of excitatory synapses, *E_i_* is the reversal potential of inhibitory synapses, and *τ_e_* and *τ_i_* are corresponding time constant. For each incoming spike, the presynaptic membrane induces an increment of postsynaptic conductance (Δ*g_e_* or Δ*g_i_*, dependent on the synapse type).

The reported simulation results were modeled with a conductance-based leaky integrate and fire model with all parameters used listed in the Tab.1.

**Table 1.** List of parameters for neurons and synapses.

| Neuron<br>parameter | Value | Description | Synapse<br>Parameter | Value | Description |
| --- | --- | --- | --- | --- | --- |
| $C_m$ | 70 pF | membrane | $E_e$ | 0 mV | excitatory |
|  |  | capacitance |  |  | reversal<br>potential |
| $V_{th}$ | -50 mV | threshold<br>potential | $E_i$ | -70 mV | inhibitory<br>reversal<br>potential |
| $V_{reset}$ | -55.8 mV | reset potential | $\Delta g_{exci}$ | 15 nS <sup>a</sup> | excitatory<br>postsynaptic<br>conductance<br>increment |
| $\tau_{ref}$ | 5 ms | refractory<br>period | $\Delta g_{inhi}$ | 10 nS <sup>b</sup> | inhibitory<br>postsynaptic<br>conductance<br>increment |
| $E_l$ | -55.9 mV | leaky reversal<br>potential | $\tau_e$ | 0.23 ms | excitatory time<br>constant |
| $g_l$ | 13 nS | leaky<br>conductance | $\tau_i$ | 2 ms | Inhibitory time<br>constant |

a. $\Delta g_{exci} = 100$ nS for the calyx of Held in the GBC-to-MNTB projection
b. $\Delta g_{inhi} = 75$ nS for the inhibitory projections to the MSO

### 2.3. Network model

The network architecture (Fig.2) of the auditory brainstem model reported here is based on our previous spiking network model of the medial superior olive (MSO) circuit [10] extended to simulate the major components of the auditory afferent pathway and auditory nuclei that facilitate ABRs. The Brian2 simulator [22] was used for the implementation of this network model. In this simulator, the integration method is handled automatically, first attempting to solve the equations exactly for linear equations and then resorting to numerical algorithms such as the Euler method when necessary. The simulation was performed with a time step of 10 µs for every calculation on the supercomputing cluster at the University of Colorado Boulder (Alpine High Performance Computing Cluster).

**Figure 2.**
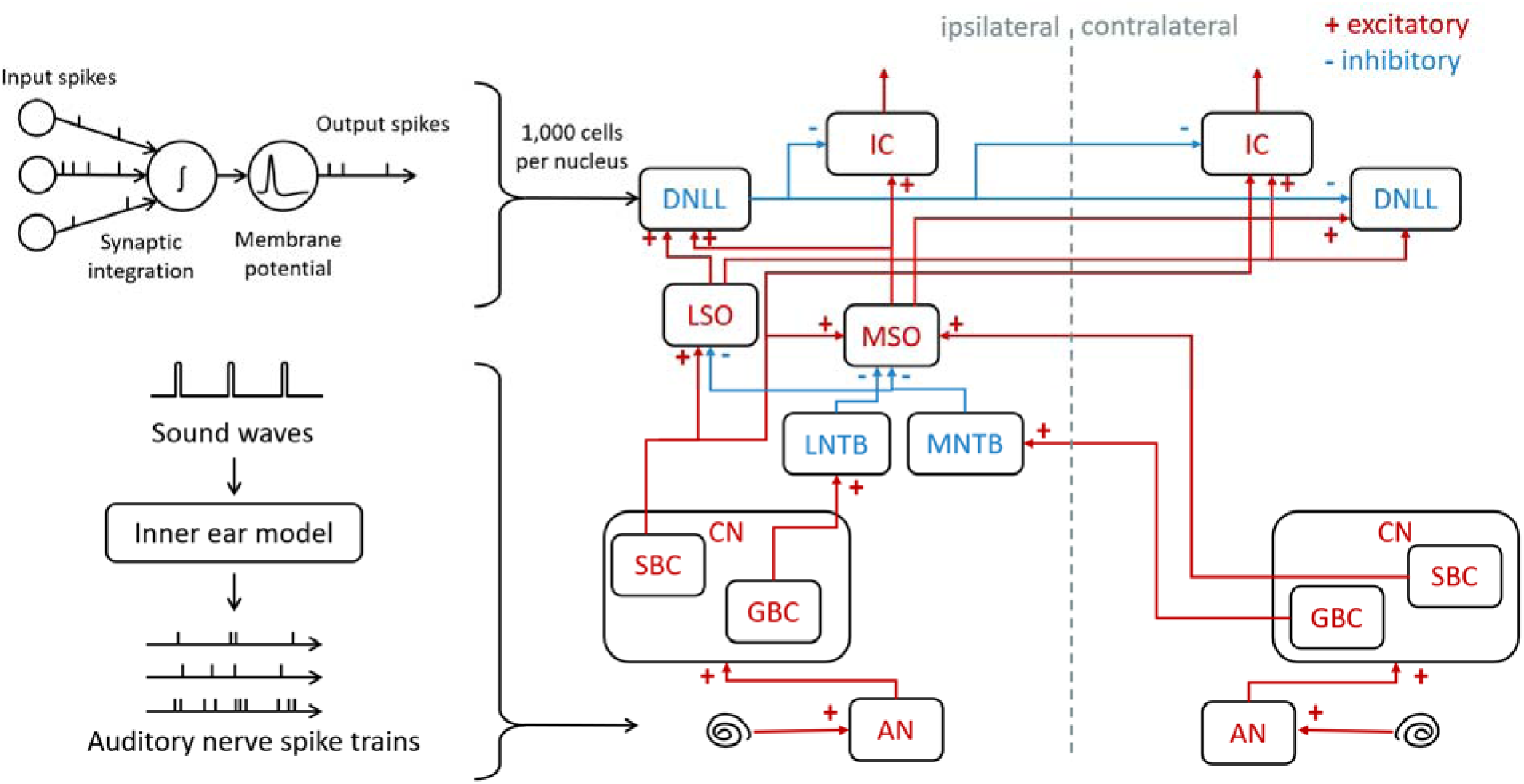
Network architecture of the auditory brainstem model.

In this model, sound waves are encoded into cochlear responses using Zilany’s model [23], implemented in the cochlear package [24], which allows for the encoding of arbitrary sound waves such as pure tones, modulated tones, chirps and natural sounds. A 100-µs complex click sound with 100 ms inter-stimulus interval was utilized for simulating click ABRs. To simulate the auditory nerve activity, we used 60% high spontaneous firing fibers, 20% medium spontaneous firing fibers and 20% low spontaneous firing fibers [25]. According to the neural circuit architecture of auditory brainstem [4], cochlear responses encoded from the auditory stimuli are sent to the brain through the auditory nerve (AN) and are received at the cochlear nuclei (CN). In the CN, the spherical bushy cells (SBCs) innervate medial superior olive (MSO) neurons bilaterally, lateral superior olive (LSO) neurons ipsilaterally, and inferior colliculus (IC) neurons contralaterally. Globular bushy cells (GBCs) innervate the contralateral medial nucleus of the trapezoid body (MNTB) and the ipsilateral lateral nucleus of the trapezoid body (LNTB). The MSO receives bilateral excitation from SBCs, contralateral inhibition via MNTB, and ipsilateral inhibition via LNTB. Meanwhile, the LSO receives ipsilateral excitation from SBC and contralateral inhibition via MNTB. Next, MSO innervates IC ipsilaterally and dorsal nuclei of lateral lemniscus (DNLL) bilaterally, and LSO innervates the ipsilateral DNLL and the contralateral IC. Finally, DNLL inhibits bilateral ICs and contralateral DNLL.

In this model, every neural population contained one thousand spiking neurons that stochastically connected to neurons in corresponding populations with a connection probability (*p_connect_*), an axonal transmission delay (*t_trans_*), and a synaptic delay (*t_syn_*). The detailed parameters of the connectivity were adopted from previous studies [10, 26–30] and are listed in Table 2. The delays in the network were randomly assigned to each synaptic connection using the mean value listed in Tab.2 with a standard deviation of 50 microseconds. The *t_trans_* of synaptic connections from contralateral side were estimated by the fiber myelin thickness and varied in the result section.

**Table 2.** List of network parameters for the auditory brainstem model.

| Population | Type | Connection | $p_{connect}$ | $\bar{t}_{trans}$ (ms) | $\bar{t}_{syn}$ (ms) |
| --- | --- | --- | --- | --- | --- |
| **AN** | Spike generator | AN → ipsi. GBC | 4 % | 0.2 | 0.6 |
| | | AN → ipsi. SBC | 4 % | 0.2 | 0.6 |
| **SBC** | Excitatory | SBC → ipsi. MSO | 0.6 % | 0.6 | 0.6 |
| | | SBC → cont. MSO | 0.6 % | 0.8 | 0.6 |
| | | SBC → ipsi. LSO | 4 % | 0.6 | 0.6 |
| | | SBC → cont. IC | 0.4 % | 0.8 | 0.6 |
| **GBC** | Excitatory | GBC → ipsi. LNTB | 0.3 % | 0.3 | 0.3 |
| | | GBC → cont. MNTB | one-to-one | 0.4 | 0.2 |
| **MNTB** | Inhibitory | MNTB → ipsi. MSO | 0.3 % | 0.1 | 0.3 |
| | | MNTB → ipsi. LSO | 0.8 % | 0.1 | 0.3 |
| **LNTB** | Inhibitory | LNTB → ipsi. MSO | 0.3 % | 0.1 | 0.3 |
| **MSO** | Excitatory | MSO → DNLL | 0.4 % | 0.3 | 0.3 |
|  |  | MSO → ipsi. IC | 0.8 % | 0.3 | 0.3 |
| **LSO** | Excitatory | LSO → DNLL | 0.4 % | 0.3 | 0.3 |
|  |  | LSO → contra. IC | 0.8 % | 0.3 | 0.3 |
| **DNLL** | Inhibitory | DNLL → contra. DNLL | 0.4 % | 0.3 | 0.3 |
|  |  | DNLL → IC | 0.2 % | 0.3 | 0.3 |
| **IC** | Excitatory |  |  |  |  |

### 2.4. Simulated ABRs

The population responses of each modeled nucleus (Fig.3a), driven by the click sound activity, were calculated separately as the peri-stimulus time histograms shown in Fig.3b. The field potentials for each nucleus were approximated using instantaneous firing rates calculated using a Gaussian filter with a kernel standard deviation of 0.05 ms on the population of spike trains from each stimulation (Fig.3c). This straightforward approach has been employed in previous ABR models to simulate individual ABR waves [11, 13]. While synaptic currents are a key contributor to field potentials, they are highly correlated with firing activity used in this approach, suggesting that this simplified method may still provide a reliable representation.

**Figure 3.**
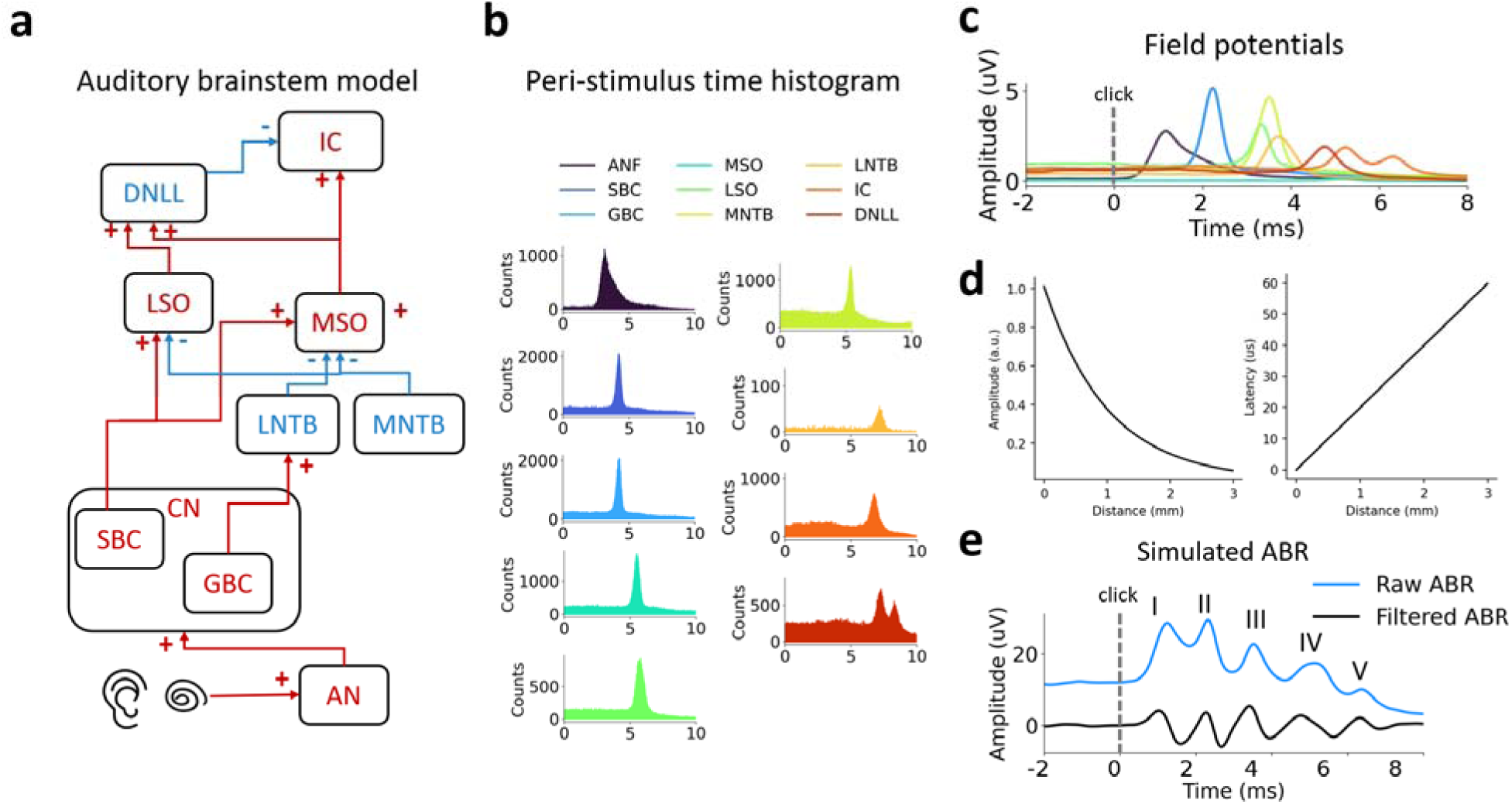
Workflow of modeling ABRs. (a) The auditory brain stem that was modeled. (b) Peri-stimulus time histograms induced by encoding click sounds averaged across all neurons for each nucleus. (c) Approximated field potentials from histograms. (d) Exponentially attenuated amplitudes and linearly increased latencies for field potentials propagating over distance. (e) Final ABRs modeled by propagating and integrating field potentials according to the geometric distribution of the nuclei and electrode location.

These field potentials were then assumed to have originated at specific spatial locations based on the geometric arrangement of these nuclei in the brain atlas for the desired species such as gerbil [31], mouse [32], or human [33]. With the location of the electrode overlayed, the Euclidean distances between nucleus centers and electrodes were calculated, and the field potentials propagated from the nuclei were estimated with amplitudes exponentially attenuating with distance, as well as the propagation latency linearly increasing with distance, assuming 50 m/s (Fig.3d). The single-trial ABR traces were then generated through integrating the field potentials of all simulated nuclei at the recording location, and the raw ABR trace was obtained by taking peri-stimulus signals averaged across all single-trial traces (Fig.3e). Finally, a detrend filter based on the Savitzky-Golay filter was applied to obtain clearer peak amplitudes and latencies, and to produce higher compatibility with existing ABR recordings from experiments.

### 2.5. ABR analysis

Experimentally recorded ABRs were analyzed with the same workflow. The raw traces were first filtered with a Butterworth bandpass filter (100-3000 Hz, 2nd order), and detrended by a Savitzky-Golay filter. Then, ABR waves were detected by searching for local maxima, and labeled sequentially. Manual inspections were performed, and adjustments made when misclassification occurred. The ABR peak thresholds were defined as 10 dB SPL below the minimum sound level with visually detectable peaks. The cross-correlation (xCorr) ABR thresholds were automatically computed based on curves fitted with sigmoids of the cross-correlations between adjacent sound levels [34]. The peak amplitude, area, latency and inter-peak intervals were measured as the peak metrics for characterizing ABR waves and BIC DN1. Statistical differences in the peak metrics were tested by using the two-sided Mann-Whiteney U test at sound levels higher than 70 dB SPL.

### 2.6. Confocal image analysis

The confocal image dataset consisted of 32 slides of the MNTB from 7 young gerbils and 26 slides from 5 old gerbils [18]. The number of MNTB cells was automatically counted with a customized script using the scikit-image library. Each 2D section of the z-stack was binarized using the Otsu thresholding algorithm [35]. Next, the binarized images were subtracted by the background obtained by the top-hat transform, and processed with a morphological erosion operation. The number of cells of the image was approximated by counting the number of contours detected in the processed image. The mean cell counts over all z-stacks were computed as the number of MNTB cells of the slide.

## 3. Results

We developed a spiking model of the auditory brainstem. This model was tested for accuracy against both mouse and gerbil data. We also tested the model’s ability to predict changes in the ABR waveforms in two conditions: Fragile-X syndrome in the *Fmr1*-KO mouse model and age-related changes to the ABR in aging wild-type gerbils. This allowed us to challenge the model but also importantly to study the underlying neural changes in aging and autism.

### 3.1. Modeling of ABRs in Fragile x syndrome

Alterations to ABRs have been reported both in children and adults with autism spectrum disorder, including prolonged ABR latencies [36] and altered peak amplitudes [3, 37]. To further investigate the neural mechanisms in the auditory pathway that might be present in autism, we used the model to simulate the ABRs in *Fmr1*-KO mice. Fragile X syndrome is the most common monogenetic form of autism spectrum disorder and it commonly causes auditory processing deficits in the auditory brain stem, as well as neurodivergent development [16, 38, 39]. Starting with the existing wild type model, we developed the *Fmr1*-KO model by tuning the parameters to match previous reports in Fragile X syndrome, including hyperexcitability [40] and myelination deficits [41]. This was done by enhancing excitatory connectivity and slowing down axonal conduction velocity, respectively. Specifically, the *Fmr1*-KO model included doubled connectivity (*p_connect_*) in the SBC-to-LSO pathway from 4% to 8% [40], and increased transmission delay ( ) from the GBC to the MNTB by 0.2 ms based on the decreased conduction velocity calculated from the change in the myelin thickness in *Fmr1*-KO mice [41] using a previously published propagation model for myelinated axons [42, 43].

Figure 4 shows click ABR and BIC responses over a range of sound intensity levels. Wave III was delayed and attenuated in the *Fmr1*-KO model and the waves IV-V were delayed when compared to the wild-type simulation (Fig.4a). Therefore, previously reported myelination deficits and hyperexcitability in the *Fmr1*-KO can alter ABR morphology. In addition, the BIC DN1 peak of *Fmr1*-KO mice was reduced (Fig.4b), delayed, or absent. Reduced BIC amplitude is associated with behavioral deficits in binaural and spatial hearing tasks in human [19] and animal models [44] of various hearing pathologies. Thus, the finding of reduced amplitude BIC here suggests a compromised sound localization ability in Fragile X syndrome. These results are also consistent with experimental findings suggesting that that *Fmr1*-KO mice showed decreased spatial hearing in behavioral tasks [16].

**Figure 4.**
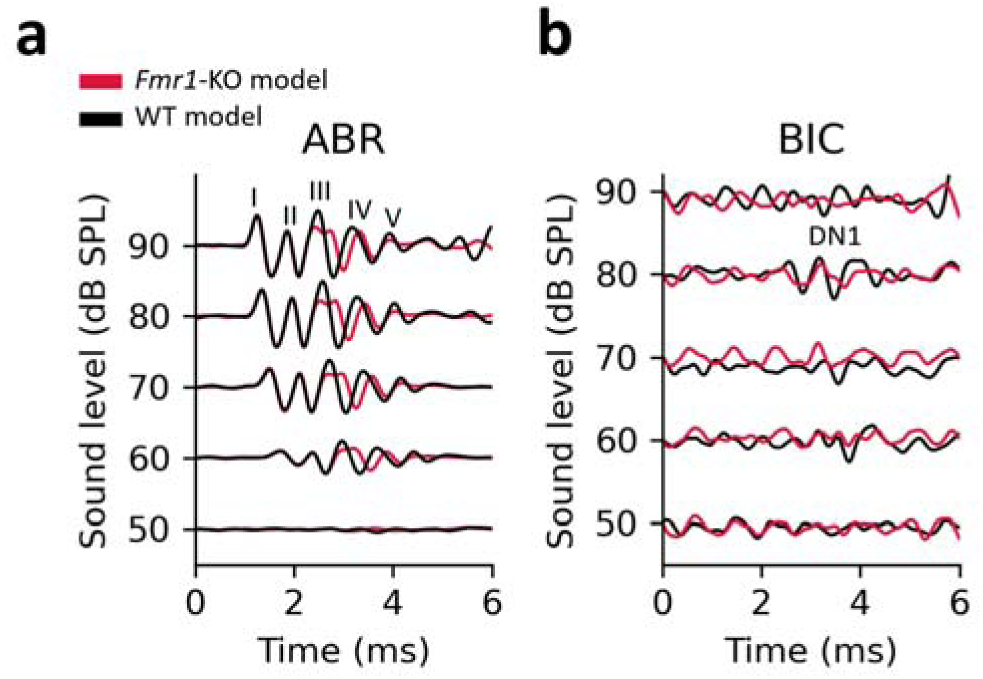
Modeled click ABRs (a) and BICs (b) in the *Fmr1*-KO (red) vs. wild type control (black) models over different sound levels. (band-pass filtered)

To verify the predictions raised by the computational ABRs, click ABR recordings were performed on Fmr1-KO and C57BL/6 wildtype mice (Fig.5). The experimentally recorded ABR traces (Fig.5a) closely matched the simulated ABRs (Fig.4a). Specifically, wave III was attenuated, and waves III-V were delayed in the Fmr1-KO condition. In addition, ABR thresholds were significantly raised by ∼10 dB SPL in the *Fmr1*-KO mice (Fig.5c) (*Fmr1*-KO: 52.61± 5.79 dB SPL (mean ± S.D.); C57BL/6: 41.32 ± 8.89 dB SPL; p < 0.0001 in Mann-Whitey U test). and BICs were significantly attenuated and delayed (Fig.5d,e).

**Figure 5.**
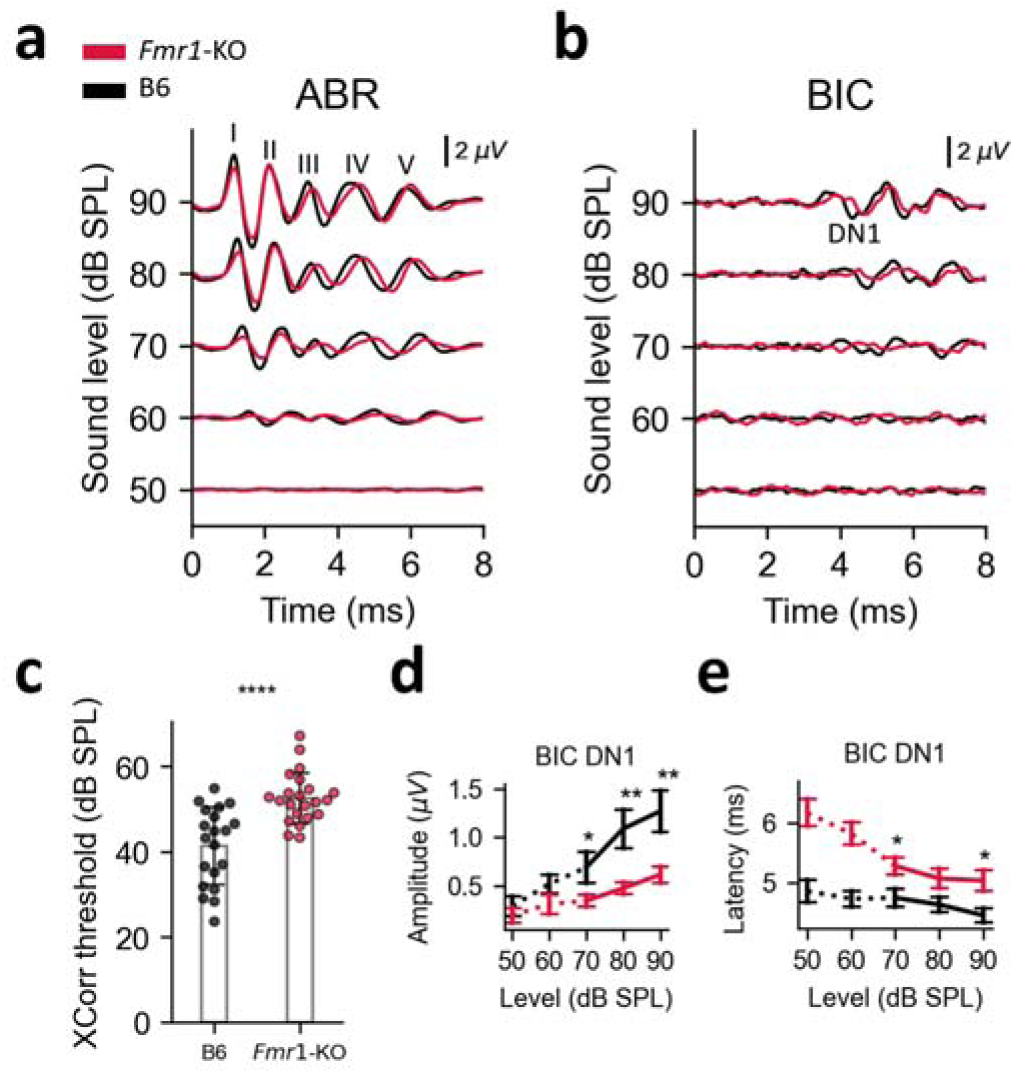
Experimentally recorded ABRs from Fmr1-KO mice (red) and C57BL/6 wildtype controls (black). (a-b) Sample ABR (a) and BIC (b) traces from wildtype and Fmr1-KO mice over different sound intensity levels. (c) Click ABR thresholds estimated with the signal cross-correlation method (Fmr1-KO: 52.61± 5.79 dB SPL (mean ± S.D.); C57BL/6: 41.32 ± 8.89 dB SPL; p < 0.0001 in Mann-Whitney U test; error bars: standard deviation). (d-e) BIC-DN1 peak amplitudes (d) and peak latencies (e) of Fmr1-KO vs. C57BL/6 mice (*: p<0.05; **: p<0.01; Mann-Whitney U test; error bars: standard error of the mean).

The statistics and reconstructed visualizations of the Fmr1-KO vs. wild type mouse ABRs are displayed in Fig.6. Despite the raised ABR threshold, the largest change was observed in wave III, which was significantly attenuated in Fmr1-KO mice (Fig.6a). Additionally, longer latencies were observed for wave III-V in KO mice (Fig.6b). The reconstructed ABR waves (Fig.6c) were calculated by fitting Gaussian functions with the averaged peak metrics. These reconstructions also show a significantly reduced wave III, and a longer latency for the following waves. These experimental results suggest that our model successfully predicts the alterations of ABR morphologies in Fragile X syndrome. Moreover, the experimentally reported myelin deficits and hyperexcitability in Fmr1-KO mice are sufficient to recapitulate the distorted ABR morphology in our model. Behaviorally, these alterations are consistent with the previously reported degradation of binaural and sound localization abilities.

**Figure 6.**
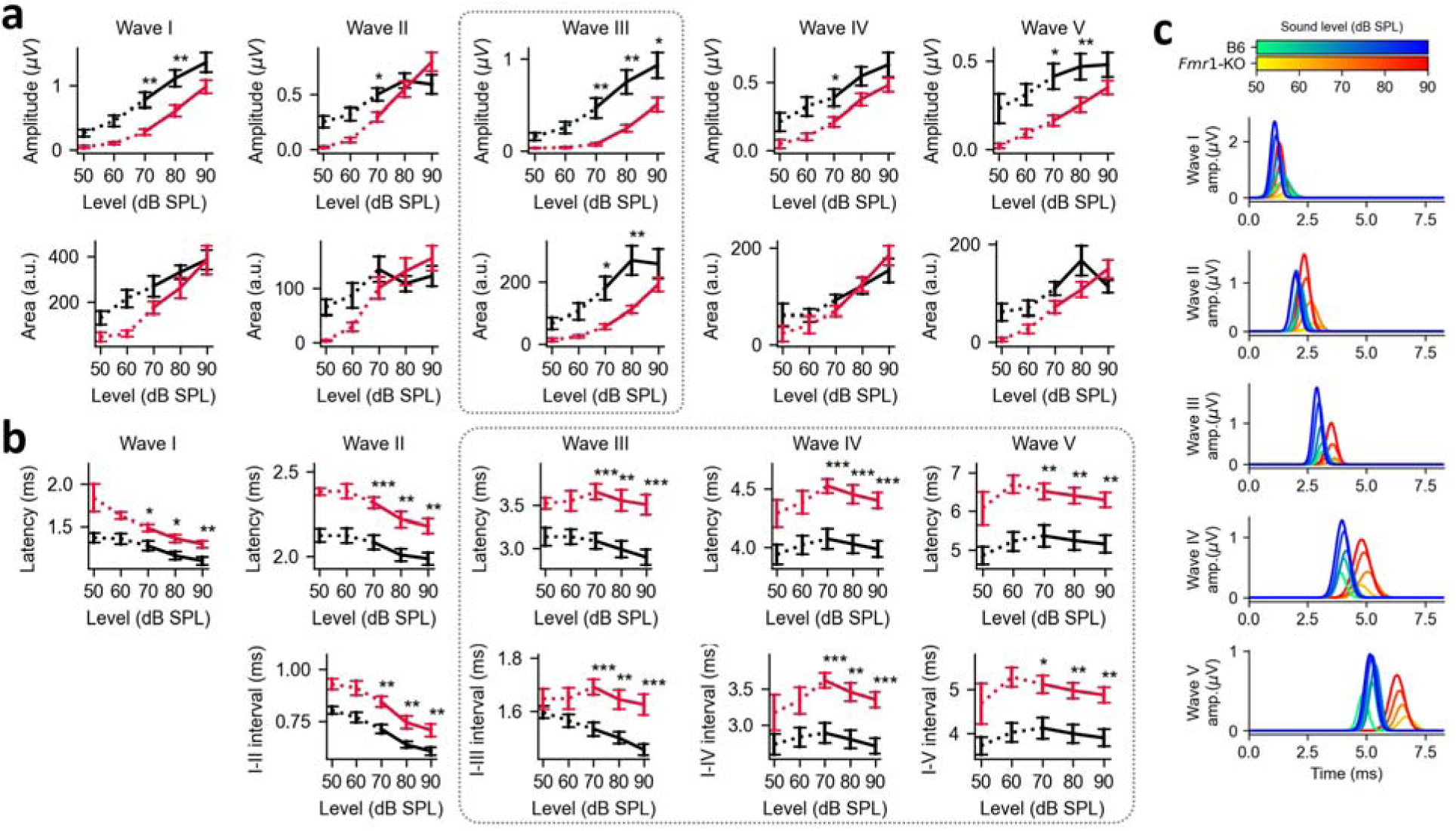
Peak metrics of experimentally measured ABR traces from *Fmr1*-KO mice vs. C57BL/6 wild type controls. (a) Peak amplitudes (top) and surface areas (bottoms). (b) Peak latencies (top) and intervals relative to wave I (bottom). (c) Overall ABR waves based on the average peak metrics from Fmr1-KO mice (orange lines) and C57BL/6 mice (blue lines). (*: p<0.05; **: p<0.01; ***: p<0.001; Mann-Whitney U test; error bar: standard error of the mean)

### 3.2. Modeling aging-related alterations in auditory brainstem

Our previous experimental study reported changes in ABR morphology in old gerbils (N=30; ages between P750 to P1167) along with alterations in myelination in the GBC-MNTB pathway and a reduction of mature oligodendrocytes in the trapezoid body when compared to young animals (N = 62; ages between P60 to P109) [18].

Briefly, in this dataset, wave I, wave III and wave IV were significantly attenuated, and the latency of wave IV was significantly shorter in old gerbils (Fig.7). As in the Fragile X model, the strongest difference was observed in wave III was the most substantially degraded wave (Fig.7e). Notably, the presence and amplitude of wave III correlates with the integrity of the MNTB or its output [45–47], suggesting altered MNTB activities in both aging and autistic animal models. These gerbil data are consistent with human studies [48], which also report a wave III decrease in older subjects. Moreover, the BIC-DN1 was decreased and had a shorter latency in old gerbils (Fig.7b). Thresholds of click ABRs did not significantly depend on age (Fig.7d; Young: 32.01 ± 12.71 dB SPL (mean ± SD); old: 29.86 ± 12.00 dB SPL; P = 0.4841, Mann Whiteney U test).

**Figure 7.**
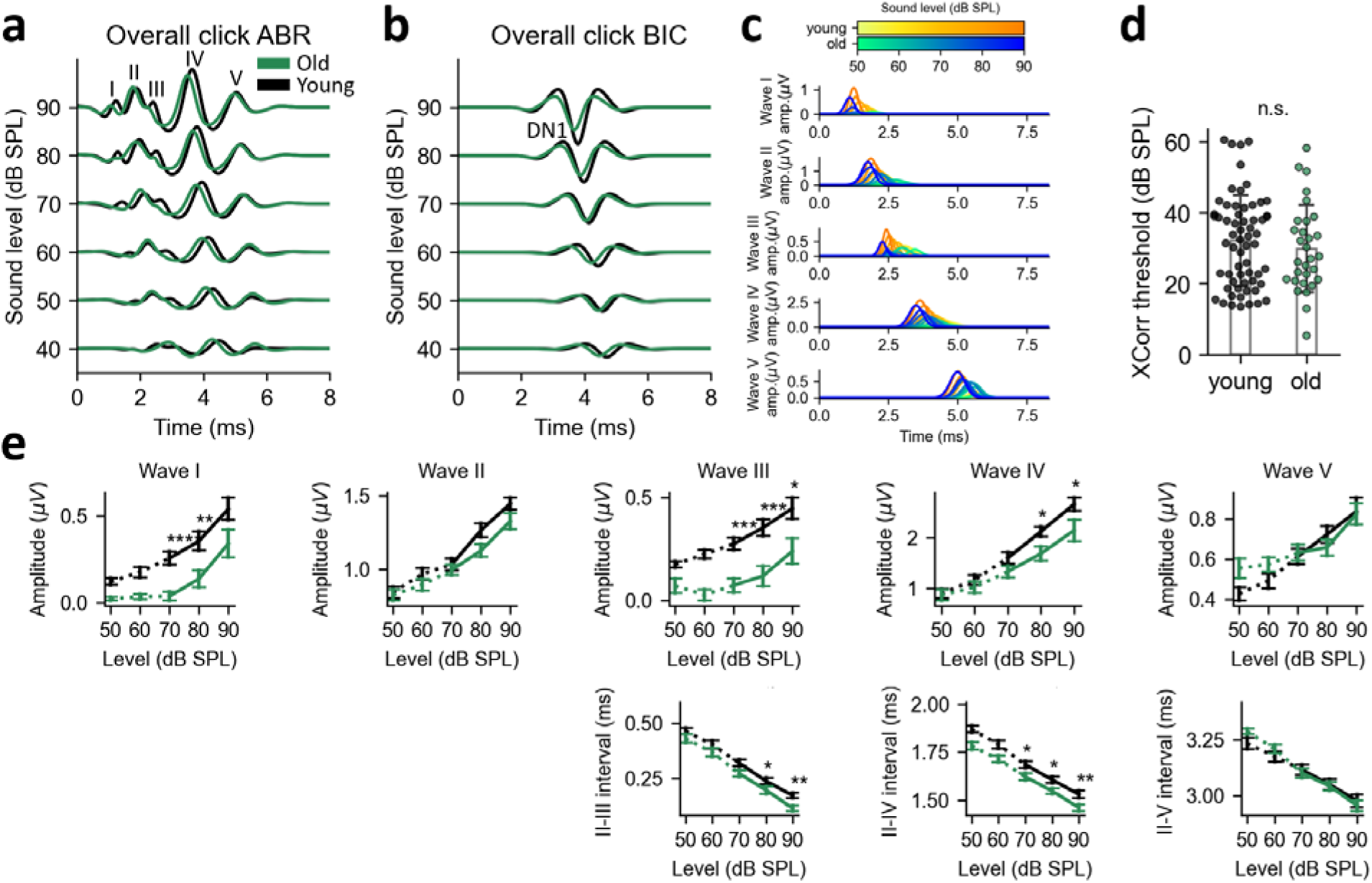
Experimentally recorded click ABRs and analysis of peak metrics in old (green) and young (black) gerbils. This is the same dataset as reported in [18]. (a-c) Overall click ABRs (a), BICs (b), and ABR waves (c) reconstructed using averaged peak metrics quantified from all experimentally recorded ABR traces. (d) Click ABR thresholds of old and young gerbils approximated with the signal cross-correlation (Young: 32.01 ± 12.71 dB SPL (mean ± SD); old: 29.86 ± 12.00 dB SPL; P = 0.4841, Mann Whiteney U test; error bar: standard deviation). (e) Peak amplitudes of ABR waves and the relative intervals from wave II. (*: p<0.05; **: p<0.01; ***: p<0.001; Mann-Whitney U test; error bars: standard error of the mean)

Based on these findings, an ABR model of aging gerbils was developed to study the causal relationship between the reported age-related changes in auditory brainstem and the altered ABR waves. Similarly to the development of the Fragile X model, several neuronal parameters were adjusted to establish this aging model. Again, the experimentally observed altered myelination leads to an increase in the modeled transmission delay ( ) from the GBC to the MNTB by 0.2 ms [18]. The excitatory postsynaptic conductance ( ) of the Calyx of Held was decreased from 100 nS to 50 nS due to maller MNTB soma size in old gerbils [49], as well as the smaller coverage ratio of the calyx of held in old rat [50]. Additionally, the number of MNTB cells was reduced from 1000 per side to 700 per side in the aging model. As all these modifications did not affect signaling at the level of cochlear nuclei, the aging model could diverge from control only in waves III-IV.

**Figure 8.**
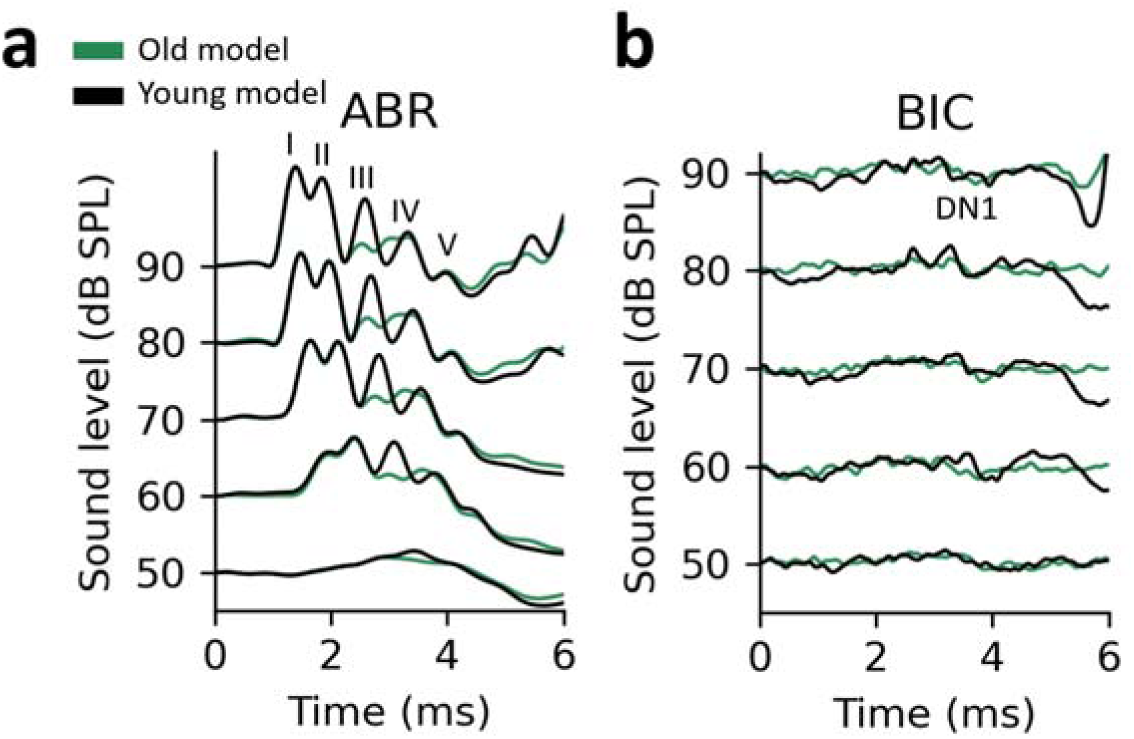
Simulated click ABRs (a) and BICs (b) in aging (green) and young (black) models over different sound levels (unfiltered raw traces).

Simulated ABRs and BICs in the aging model present a trend with smaller or absent wave III and earlier onset of wave IV, consistent with the experimentally recorded ABR changes in old gerbils. Therefore, the altered ABR wav III and wave IV in old gerbils could be explained by the demyelination and degradation along the contralateral inhibitory pathway in the superior olivary complex.

This particular aging model contains an assumption that the number of MNTB cells is reduced in aged gerbils. Although a reduction in the MNTB cell count was reported in old C57BL/6 mice [51], this strain exhibits severe hearing loss as early as 6 months, begging the questions whether this phenomenon also occurs in other strains and species [52, 53]. To further explore this point, MNTB cell counts were performed from confocal images in old gerbils. In these confocal images, MNTB neurons were sparser in old gerbils than in young animals (Fig.9a). The estimated cell counts were significantly different with relatively fewer cells in sections from old gerbils (Fig.9b; Ol : 113.49 ± 18.90 cells, N_animal_ = 5, n_slides_ = 26; Young: 133.33 ± 34.51 cells, N_animal_ = 7, n_slides_ = 32; p = 0.0038, Mann-Whiteney U test). These cell counts further confirm the age-dependent degradation of the contralaterally evoked inhibition by MNTB in old gerbils, exactly as postulated by the computational model.

**Figure 9.**
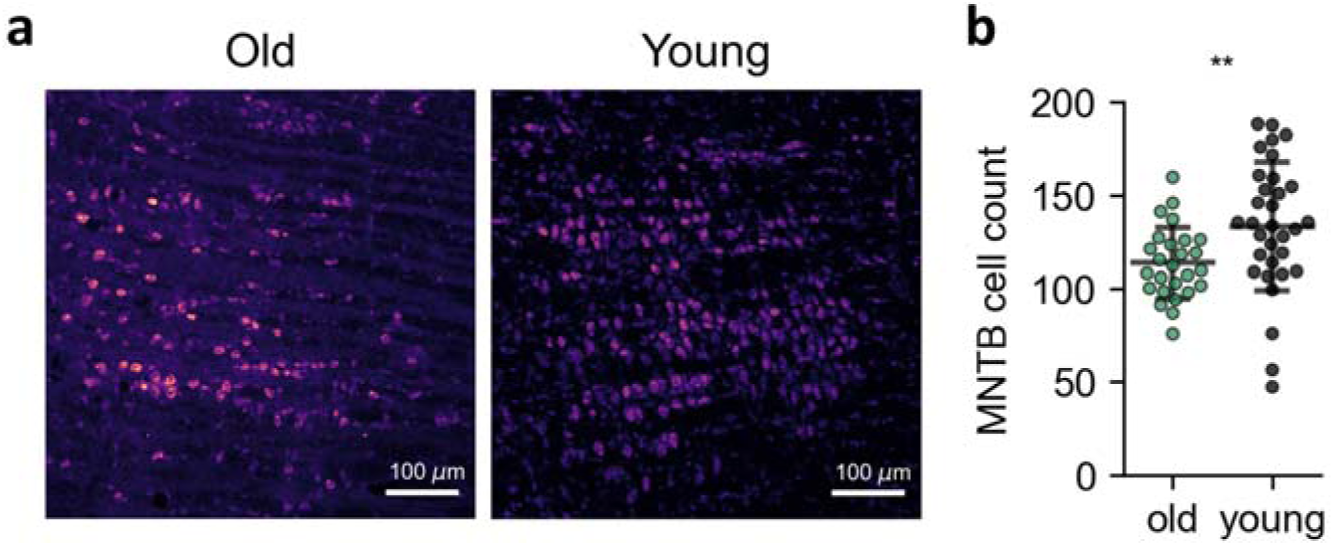
Quantifying MNTB cells in old and young gerbils. (a) Confocal images of MNTB sections from old gerbils (left) and young gerbils (right). (error bar: 100 um) (b) Estimated number of MNTB cells per 640 um * 640 um slide. (Old: 113.49 ± 18.90 cells, N_animal_ = 5, n_slides_ = 26; Young: 133.33 ± 34.51 cells, N_animal_ = 7, n_slides_ = 32; p = 0.0038, Mann-Whiteney U test; error bar: standard deviation)

## 4. Discussion

There are several results of this study: First, we were able to extend an integrate and fire model to a population level such that we could predict ABRs correctly by modeling the neuronal populations of auditory brain stem nuclei involved in the sound localization process. While ABR models have been proposed before, these did either only model single waves of the ABR trace, or lacked the single-cell approach we used here. The model’s ability to simulate the ABR trace from wave I to wave V based on neuronal activities provides a considerable advancement in understanding auditory processing and its associated dysfunctions.

Second, our model not only predicted ABRs of the healthy auditory system correctly, but also correctly captured neurological alterations which are present in two conditions, namely autism and age-related central hearing loss. Both of these conditions have in common that affected listeners have trouble isolating a sound source of interest among competing background noises. In such environments, commonly termed “cocktail-party situations”, listeners use their sound localization circuit to spatially segregate sound sources of interest from background sounds based on spatial location. Human listeners with normal hearing are able to localize to 5 spatial degrees or better, such that competing sounds would fall into separate spatial channels as long as they are at least 5 degrees removed from each other. Any alterations in the sound localization pathway which results in less precise spatial location should then lead to difficulties functioning in cocktail party environments. Both listeners with autism spectrum disorder and aging listeners are well known to have difficulties in acoustically busy environments, and in both conditions distinct alterations in the sound localization pathway have been demonstrated across species. These alterations were recapitulated by our model based on reductions in the amplitude of the BIC of the ABR.

Third, the results of this study underscore the utility of computational models in bridging the gap between experimental and computational work. In this particular study, the model predicted a decrease in the number of MNTB principal neurons in aging, which prompted a follow up experimental study with the goal to test whether this modeling prediction could be verified in anatomical analysis – the prediction was indeed verified. In our opinion, an ideal modeling approach should be both strongly based on experimental data, but should also able to point out potential gaps in the experimental data, and suggest distinct and precise follow up bench experiments, as it happened in this case.

Specifically, the application of the model in Fragile X syndrome recapitulated alterations of the ABR morphology driven by myelin deficits and hyperexcitability, such as decreased wave III amplitudes and prolonged wave III-V intervals. These findings align with previously reported results in children and adults with autism [37, 54–56]. The validation of these simulated results with experimentally recorded ABRs from Fmr1-KO mice suggests that the myelin deficits and hyperexcitability are sufficient to cause the observed ABR alterations and are also consistent with auditory phenotypes in autism. These findings support potential therapeutical approaches through remyelination [57] or by modulating neuronal connectivity [58].

Regarding the aging phenotype, our model successfully approximated the ABRs recorded from aging gerbils, revealing deficits in the contralateral inhibitory pathway and a decrease in the cell count in MNTB, which was subsequently validated in confocal images. These results confirm a prior study showing reduced wave I, III, IV and BIC in aged gerbils [59]. Again, these findings suggest therapeutic approaches in restoring this form of age-related hearing loss in elderly individuals through remyelination [60] and by enhancing contralateral inhibition. We note that while contralateral inhibition has been investigated in many studies, the ipsilateral inhibition via LNTB is much less understood, and we do not know if this projection also altered in autism or aging. Because of the paucity of experimental data, we were unable to assess potential roles of this projection in Fragile X or aging in the model.

The results presented here were obtained by using relatively simple spiking neuron models with homogeneous configurations. Future work could focus on integrating more detailed neural models with heterogenous cell types. While principal cells in each one of the nuclei modeled here are very similar in their physiological properties, differences in these properties have been observed between the nuclei. Additionally, challenging the model with more complex sound stimuli and binaural cues [61] or speech [62] would be logical next steps. This would enhance the model’s accuracy in simulating complex auditory processing tasks and disorders. Additionally, the model’s application could be extended to other neurological conditions that affect auditory processing and brain stem functions, such as schizophrenia [63], multiple sclerosis [64], or brain injury [65].

In summary, the proposed computational model of the auditory brainstem developed in this study serves as a useful tool for linking ABR recordings to the underlying neural mechanisms. The validation of the model in both autistic and aging animal models suggests its potential for various applications in auditory research and diagnostics. Future advancements in this modeling approach may provide further insights into the complexities of auditory processing.

## Acknowledgements

This work utilized the Alpine High Performance Computing Resource at the University of Colorado Boulder. Alpine is jointly funded by the University of Colorado Boulder, the University of Colorado Anschutz, Colorado State University, and the National Science Foundation (award 2201538).

## Author Contribution

B.-Z. L.: Conceptualization, Formal analysis, Investigation, Methodology, Software, Validation, Visualization, Writing – original draft. S. P.: Data curation, Investigation, Validation, Writing – original draft. M. R.: Investigation, Writing – original draft, Writing – review and editing. D. T.: Conceptualization, Writing – review and editing. T. L.: Conceptualization, Software, Supervision, Writing – review and editing. A. K.: Conceptualization, Funding acquisition, Project administration, Resources, Supervision, Writing – review and editing.

## Code Availability

The implementation source code of the presented model will be available on a public repository after this manuscript is accepted. For the peer review, the code is available on Github repository (https://github.com/libenzheng/Auditory-Brainstem-Response-Model) accessible using git clone command with the following personal access token: (github_pat_11AKAILHI0jZZcx8VO93lp_Blr5vQopLXwhlKomUm1zSBsv0aN8fD96JrT5SaPwLEA6PDQIFMHJ C44VXlY).

## Data Availability

Electrophysiology data are available from the corresponding author upon reasonable request.

## Statements and Declarations

### Ethics Declarations

All experimental procedures were reviewed and approved by the institutional animal care and use committees (IACUC) of the University of Colorado Anschutz Medical Campus (permit no.00617) and complied with all National Institutes of Health (NIH) and OLAW guidelines for the humane treatment of laboratory animals.

